## Supplemental Figures and Text for "Meta-Research: understudied genes are lost in a leaky pipeline between genome-wide assays and reporting of results"

**Supplementary files:**

- Figure S1-S12 (*this document*)
- Movie S1 (movie_s1.mp4)
- Tables S1-S5 (.xlsx *files*)
- Table S6 (table_s6.csv)

**Figure S1:** PRISMA diagram for the selection of genome-wide association studies (GWAS, from studies indexed by the NHGRI-EBI GWAS catalog^1^).

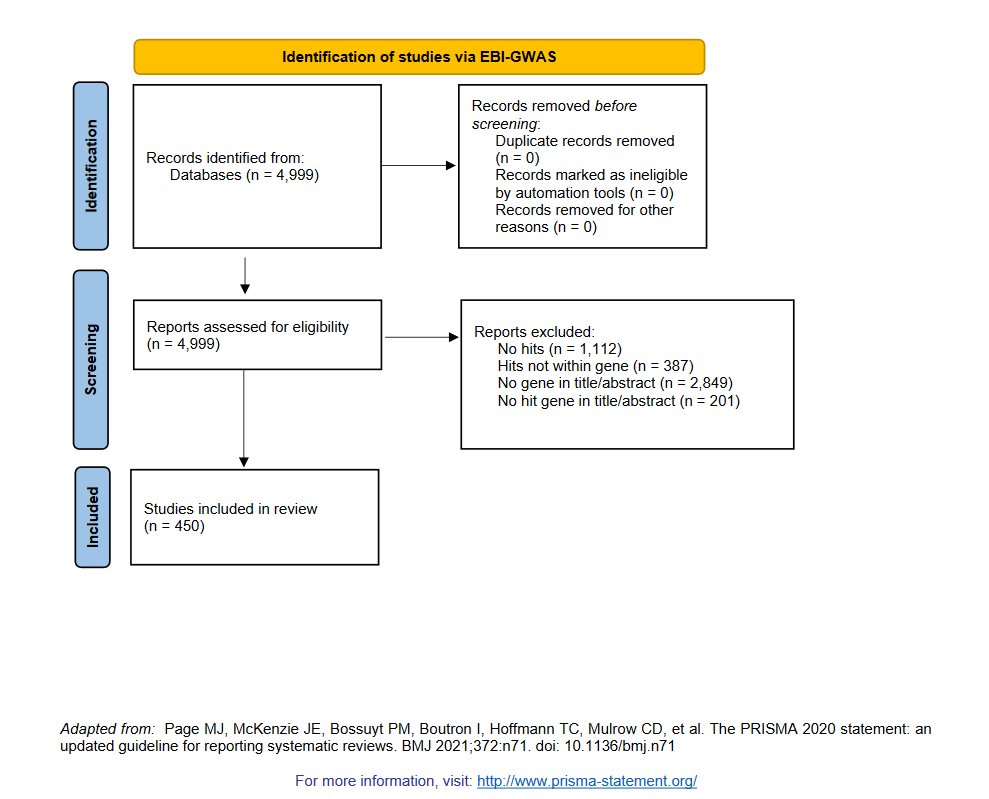

**Figure S2:** PRISMA diagram for the selection of affinity purification–mass spectrometry (AP-MS, indexed by BioGRID^2^).

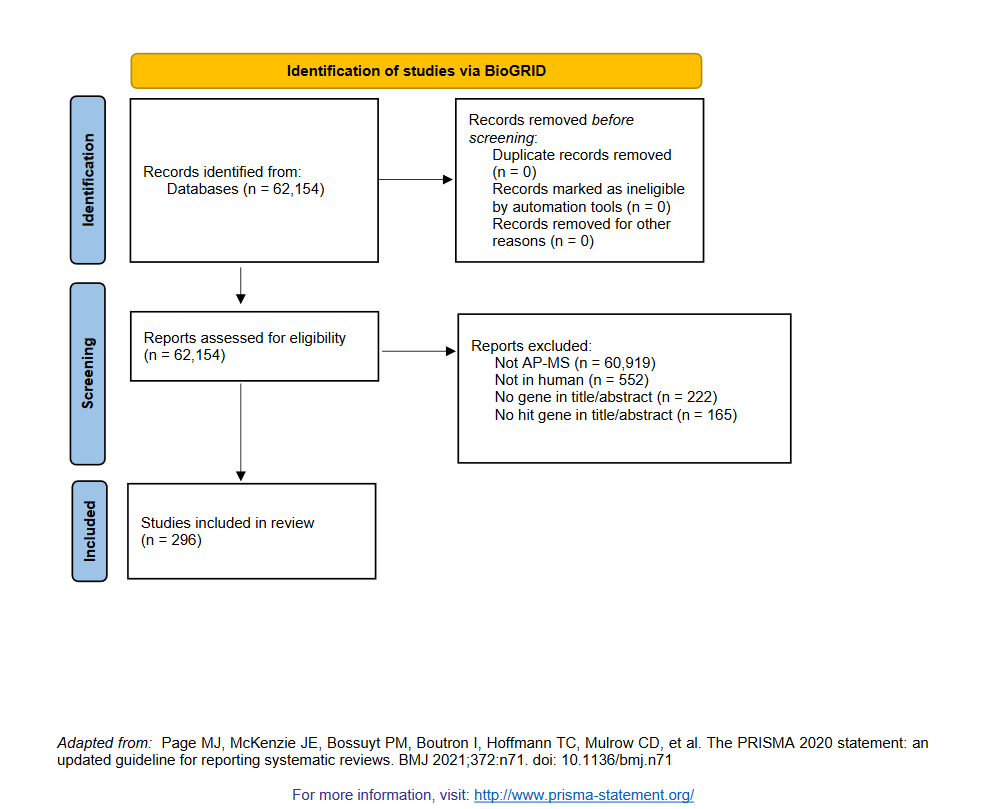

**Figure S3:** PRISMA diagram for the selection of transcriptomic studies (indexed by the EBI Gene Expression Atlas^3^).

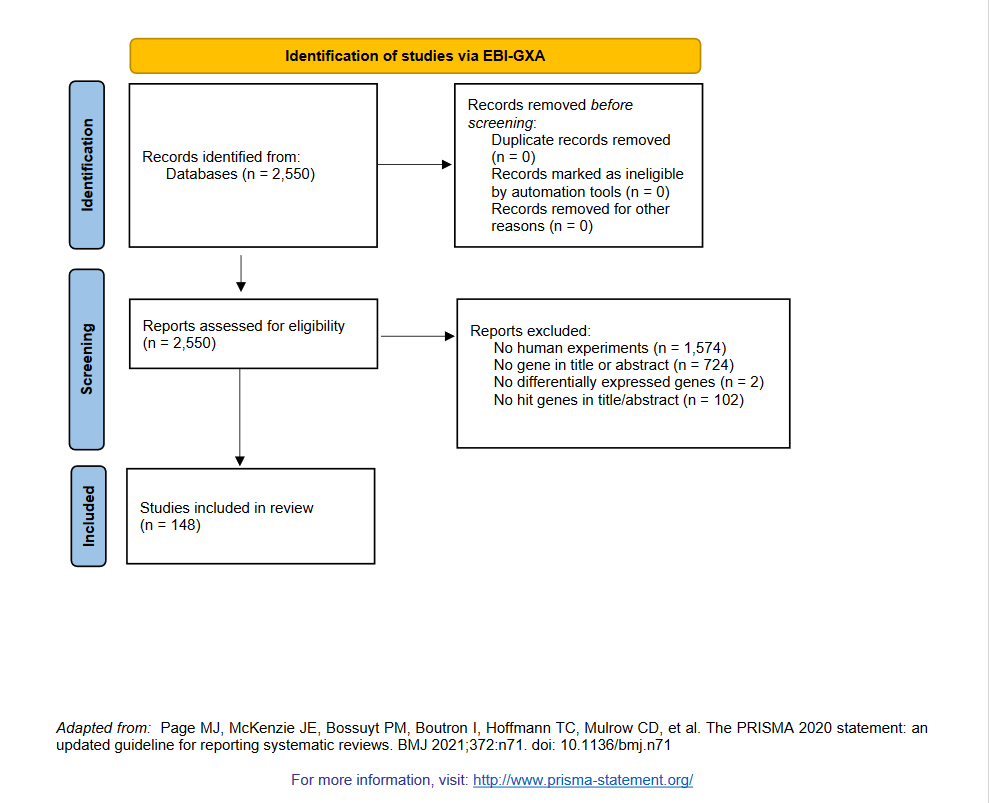

**Figure S4:** PRISMA diagram for the selection of genome-wide screens using CRISPR (indexed by BioGRID Open Repository of CRISPR Screens^2^).

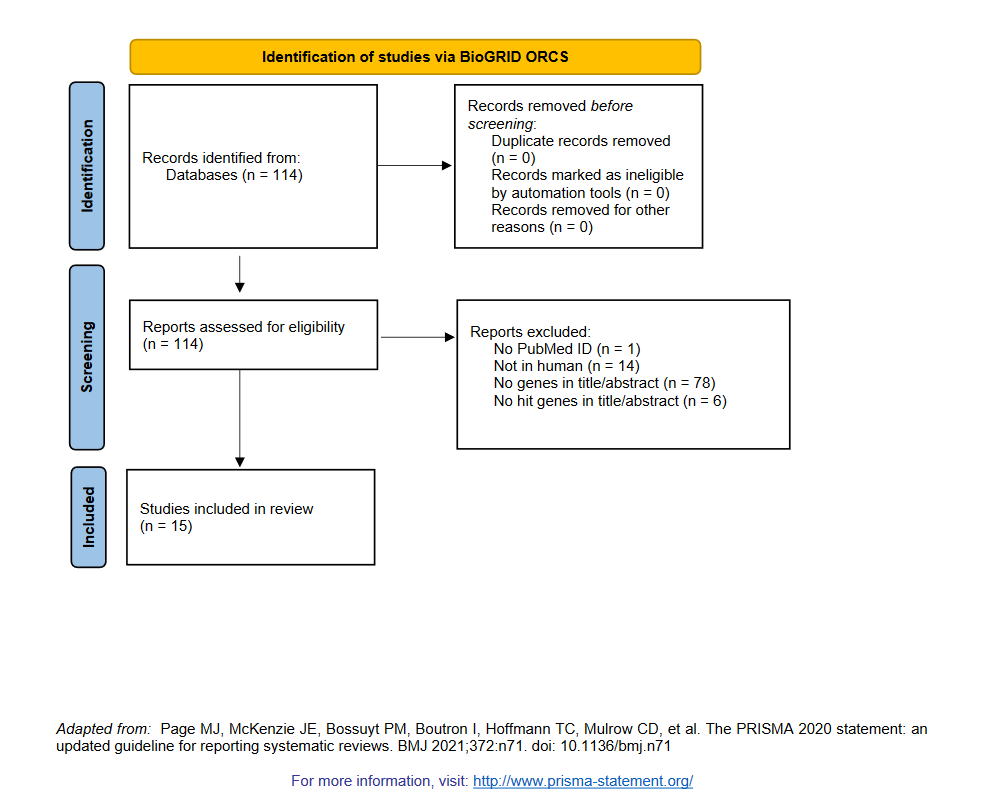

**Figure S5:** Variant of **Figure 1B** with alternate p-value/FDR thresholds for significance where applicable. GWAS p-value threshold for significance was set to 1e-10 and transcriptomics FDR threshold for significance was set to 0.0001. Alternative thresholding was not available for AP-MS or CRISPR because significance calling was complex and varied across experiments in both categories. *** denotes p < 0.001 by two-sided Mann-Whitney U test.

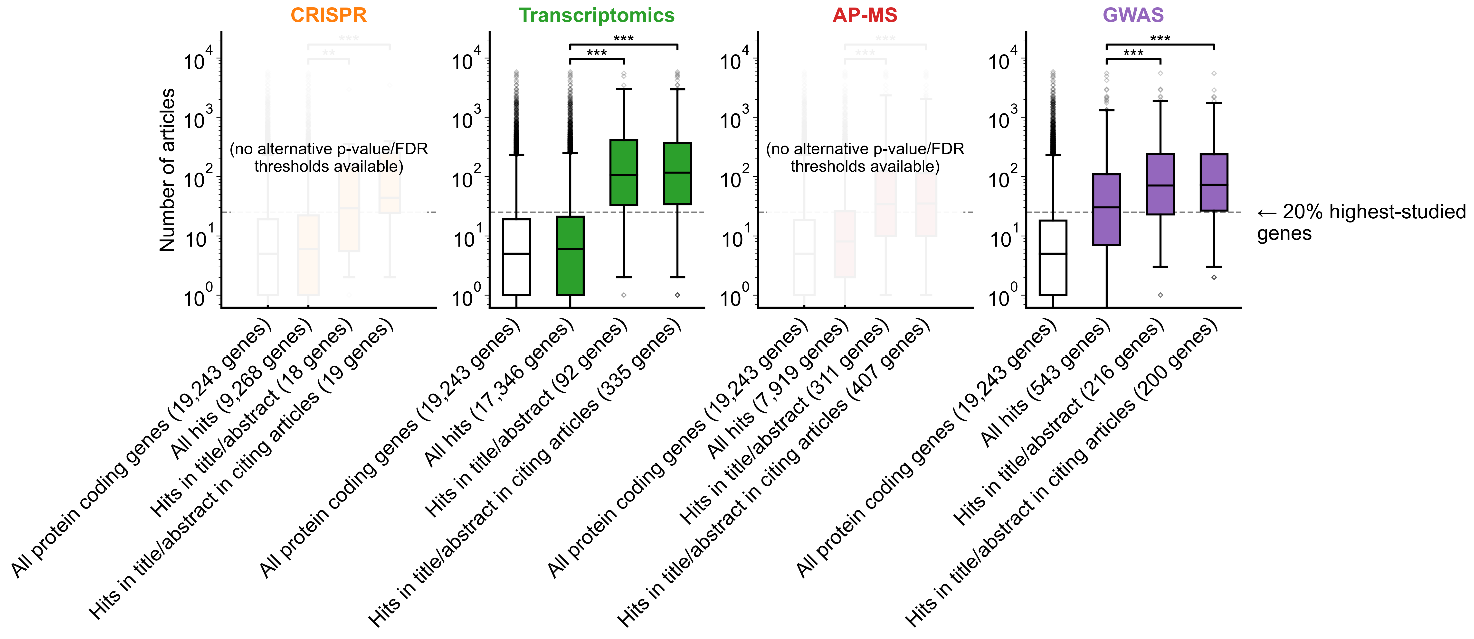

**Figure S6:** Variant of **Figure 1B** only considering articles published in 2002 or before, prior to the publication of any of the articles featuring -omics experiments which we considered for this analysis. *** denotes p < 0.001 by two-sided Mann-Whitney U test.

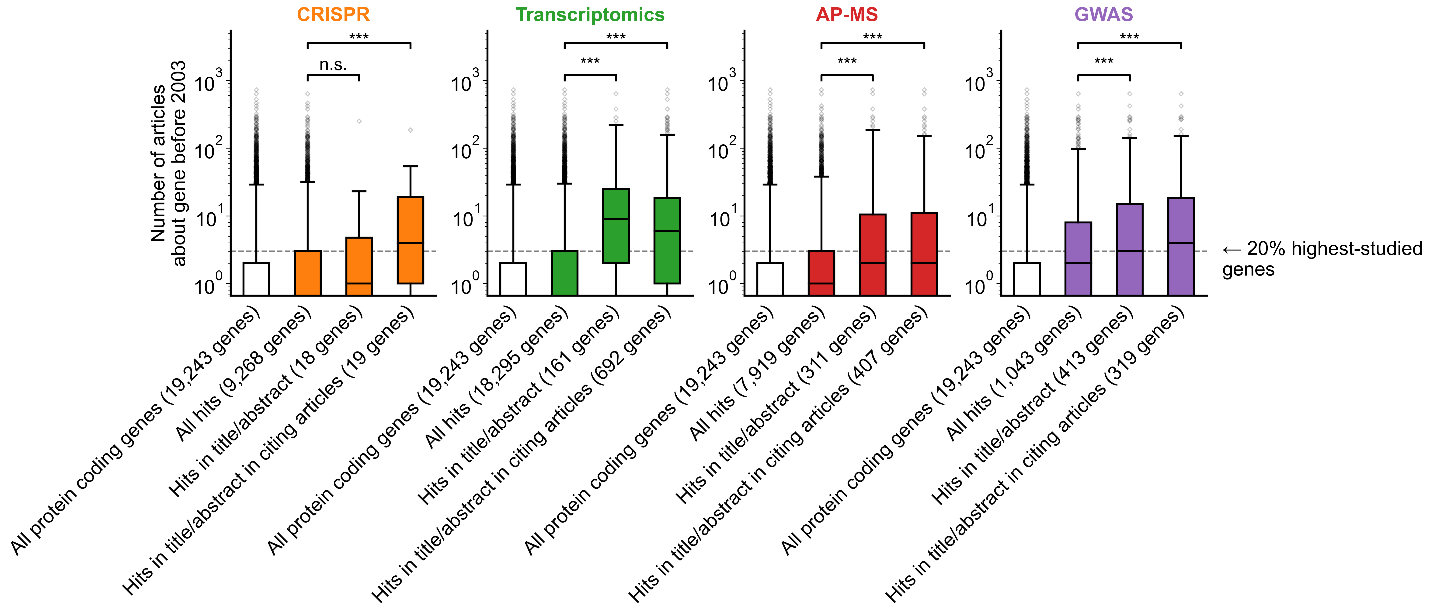

**Figure S7:** Variant of **Figure 1B** only considering one randomly chosen gene per article title/abstract. * denotes p < 0.05 and *** denotes p < 0.001 by two-sided Mann-Whitney U test.

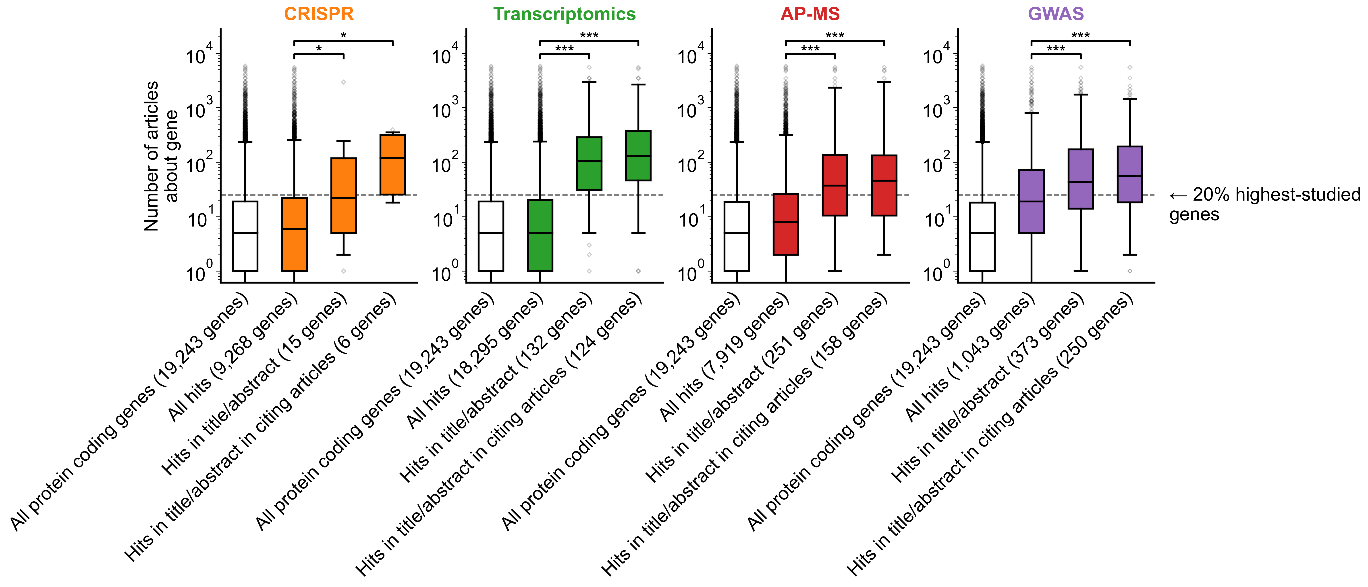

**Figure S8:** Likelihoods of being highly cited (top 5% of citations among all articles about genes, panel **a**) or lowly cited (bottom 5% of citations among all articles about genes, panel **b**) for articles about the most popular genes (top 5% accumulated articles) versus articles about the least popular genes (bottom 5% accumulated articles) by year of publication. Only articles with a single gene in the title/abstract are considered. Shaded regions show ±1 standard error of the proportion.

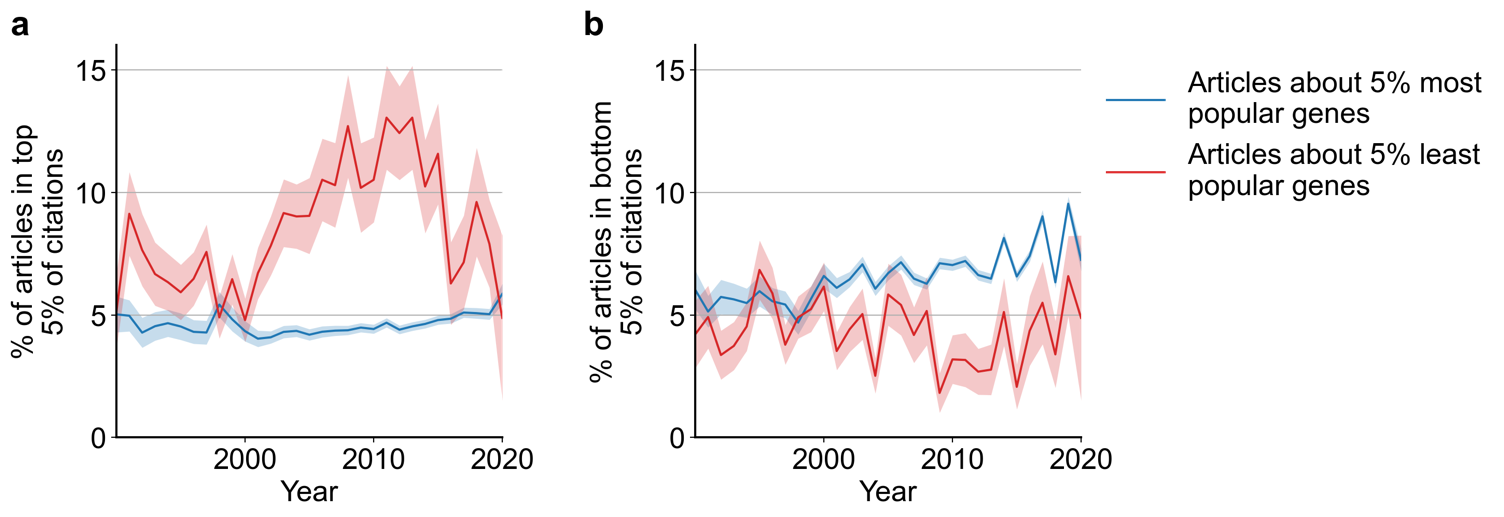

**Figure S9:** Spearman correlation and significance^4,5^ of normalized gene popularity vs normalized article citation rank for articles within disease MeSH terms from 2014 to 2018. Only articles with a single gene in the title/abstract are included. **a,** volcano plot of Spearman correlations, where size of dot corresponds to number of articles meeting criteria for each MeSH term. **b-g,** plots for several MeSH terms, corresponding to letters highlighted in **a.** Solid red lines show LOWESS regression. Only MeSH terms under the ‘Diseases’ heading (tree number starting with ‘C’) with 30 articles or more were considered (602 MeSH terms).

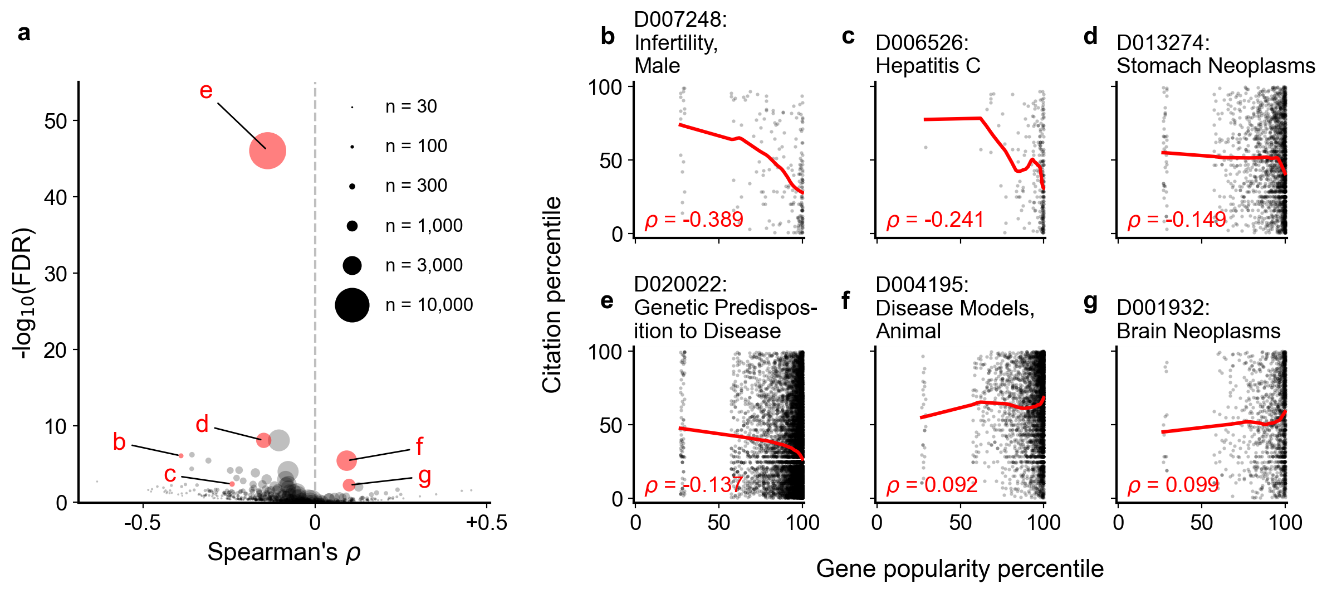

**Figure S10:** Spearman correlation and significance^4,5^ of normalized gene popularity vs normalized article citation rank for articles within technique-related MeSH terms from 2014 to 2018. Only articles with a single gene in the title/abstract are included. **a,** volcano plot of Spearman correlations, where size of dot corresponds to number of articles meeting criteria for each MeSH term. **b-g,** plots for several MeSH terms, corresponding to letters highlighted in **a.** Solid red lines show LOWESS regression. Only MeSH terms under the ‘Investigative Techniques’ heading (tree number starting with ‘E05’) with 30 articles or more were considered (264 MeSH terms).

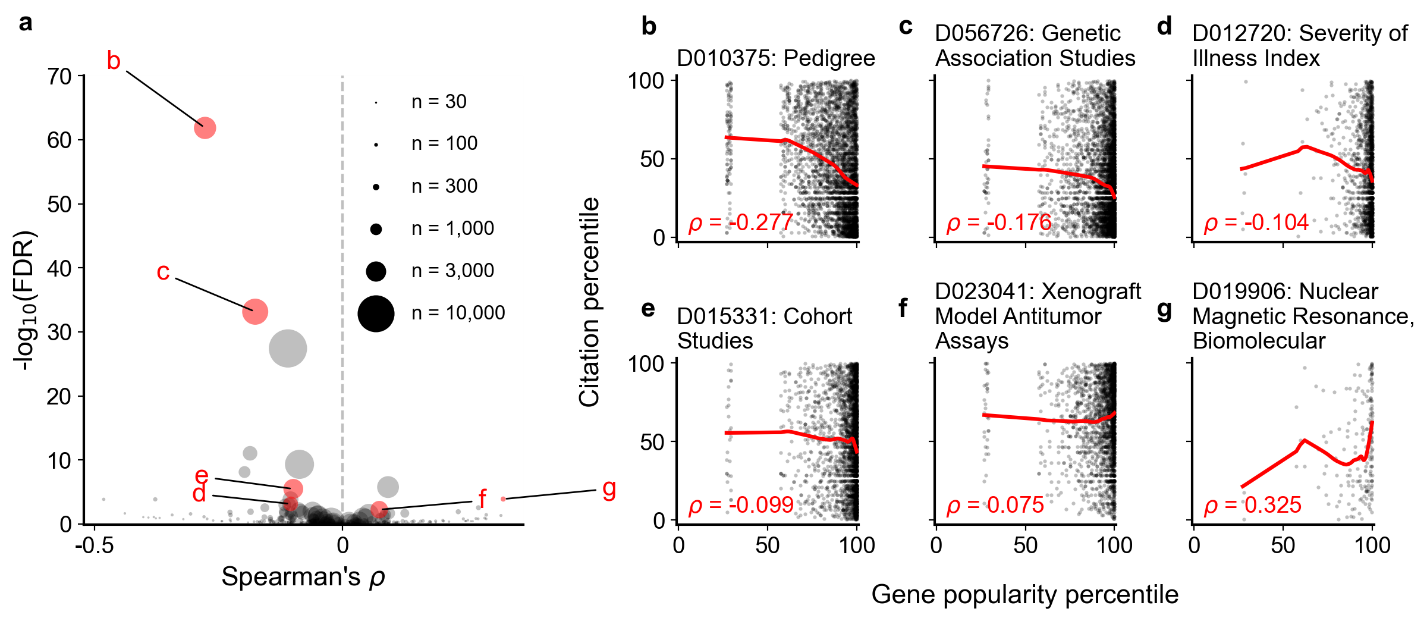

**Figure S11: Identification of default factors in FMUG, manually chosen to represent different clusters of factors.** Factors are shown along the x axis, with genes along the y axis. Binary factors are coded to 0 (white) and 1 (purple), while continuous factors are ranked from 0 to 1 with ties resolved to minimum rank. Clustering was performed with Ward’s method for hierarchical clustering^6^. Bold indicates FMUG ‘s default factors, which we selected based on this clustering and based on their strength of association with gene selection (**Figure 3, Table S2** and **Table S3**).

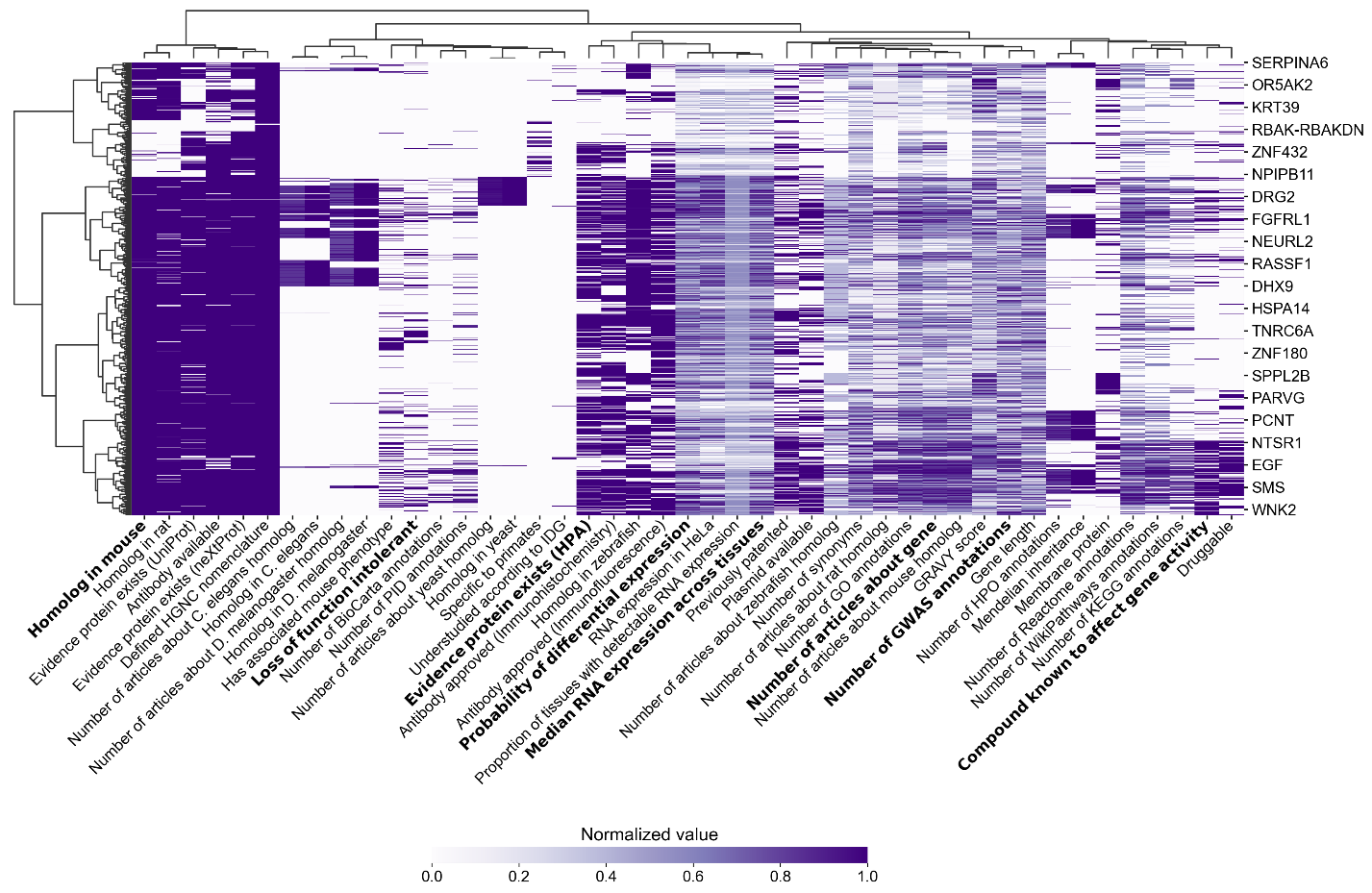

**Figure S12:** Similarity matrix and clustermap showing Spearman correlation between factors across all human protein-coding genes. Correlations involving binary factors should not be interpreted as statistical fact, as Spearman’s ρ does not account for non-ordinal data. Clustering was performed with Ward’s method for hierarchical clustering^6^.

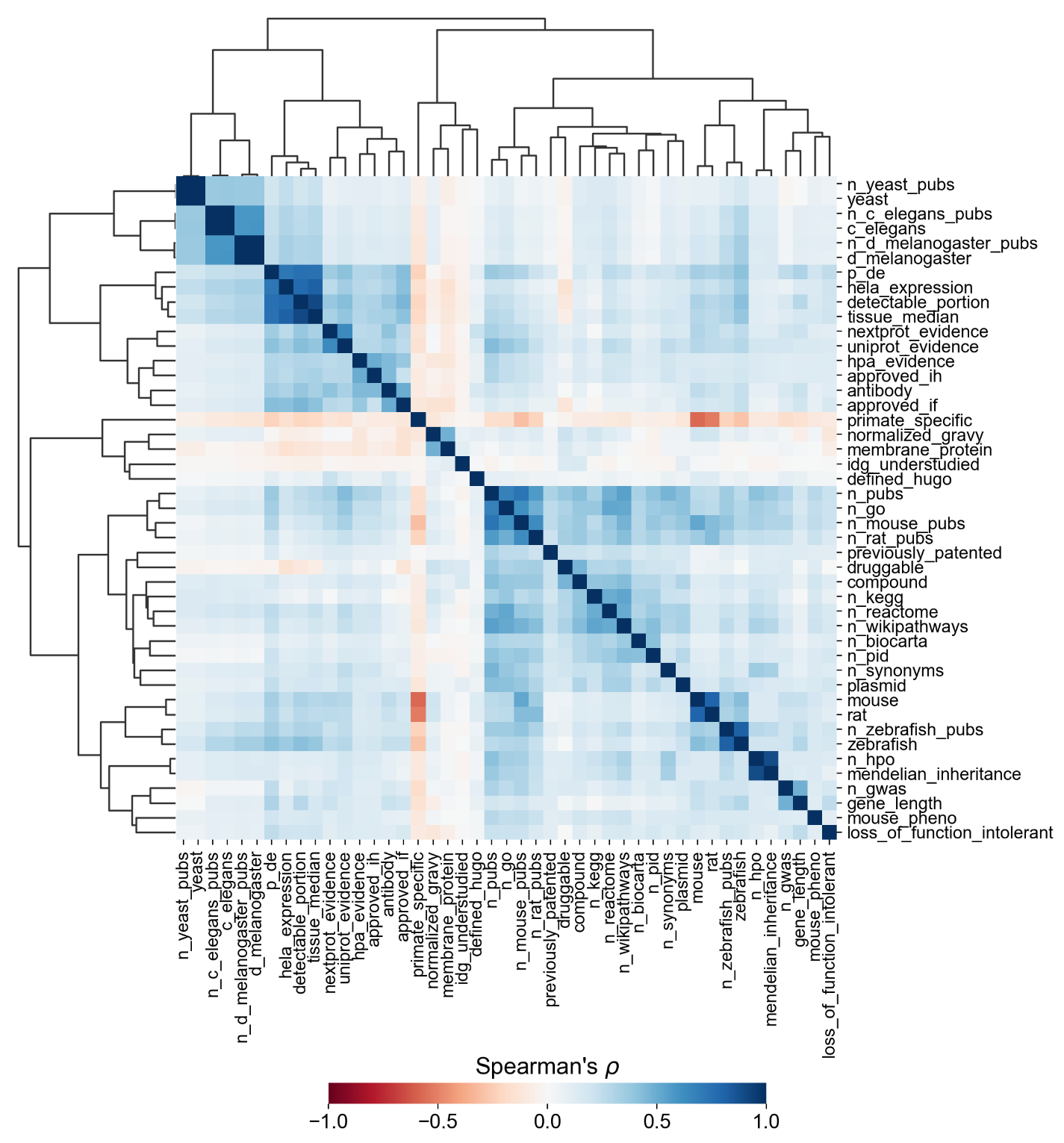

**Table S1 (*table_s1.xlsx*):** Collected factors from literature search.

**Table S2 (*table_s2.xlsx*):** Association between factors with binary (True/False) identities and highlighting hits in title/abstract of reporting articles.

**Table S3 (*table_s3.xlsx*):** Association with factors with continuous identities and highlighting hits in title/abstract of reporting articles.

**Table S4 (*table_s4.xlsx*):** Disease-related MeSH terms with a significant association between gene popularity and citations.

**Table S5 (*table_s5.xlsx*):** Technique-related MeSH terms with a significant association between gene popularity and citations.

**Table S6 (*table_s6.csv*):** PubMed ID’s and mentioned genes for collected GWAS, CRISPR, transcriptomics and AP-MS articles.

**Movie S1 (*movie_s1.mp4*):** FMUG tutorial video.
